# Planktonic Protist Biodiversity and Biogeography in Lakes From Four Brazilian River-Floodplain Systems

**DOI:** 10.1101/337824

**Authors:** Guillaume Lentendu, Paulo Roberto Bressan Buosi, Adalgisa Fernada Cabral, Bianca Trevisan Segóvia, Bianca Ramos de Meira, Fernando Miranda Lansac-Tôha, Luiz Felipe Machado Velho, Camila Ritter, Micah Dunthorn

## Abstract

While the biodiversity and biogeography of protists inhabiting many ecosystems have been intensely studied using different sequencing approaches, tropical ecosystems are relatively under-studied. Here we sampled planktonic waters from 32 lakes associated with four different river-floodplains systems in Brazil, and sequenced the DNA using a metabarcoding approach with general eukaryotic primers. The lakes were dominated by the largely free-living Discoba (mostly the Euglenida) and Ciliophora unlike previously sampled Neotropical environments, but the community similarities between samples were likewise low. These protists inhabiting these floodplains potentially form part of the large diversity of unknown diversity in the tropics.

---

Floodplains are one of the major Neotropical ecosystems, which form a fluctuating transition between aquatic and terrestrial areas due to seasonal flooding (Junk et al. 2014). About 50% of the floodplains in South America are located in Brazil, which covers 20% of the country (Junk et al. 2014; Naranjo 1995). These Brazilian wetlands contribute significantly to the overall biodiversity of South American and they have been extensively studied taxonomically for animals and plants (e.g., Desbiez et al. 2010; Gomes et al. 2012; Gopal and Junk 2000; Haugaasen and Peres 2006; Machado et al. 2015; Sanchez et al. 1999). By contrast, little is know about the microbial eukaryotic inhabitants of Brazilian river floodplains, except for some studies that focused on individual taxa (e.g., Arrieira et al. 2016; Buosi et al. 2014; Lansac-Tôha et al. 2016; Rejas et al. 2005; Schwind et al. 2017; Segovia et al. 2017).

A powerful approach to uncovering the biodiversity and biogeography of protists inhabiting any ecosystem is to use metabarcoding, where primers amplify specific DNA regions that are then sequenced on a massive scale (Creer et al. 2016; Taberlet et al. 2018). Most protist metabarcoding studies have analyzed marine near-shore and open waters, and temperate freshwaters and soils (e.g., Aguilar et al. 2016; de Vargas et al. 2015; del Campo et al. 2015; Egge et al. 2015; Filker et al. 2016; Grossmann et al. 2016; Khomich et al. 2017; Lentendu et al. 2014; Logares et al. 2014; Massana et al. 2015; Monier et al. 2015; Stoeck et al. 2010; Venter et al. 2017). Fewer metabarcoding studies have analyzed tropical ecosystems (e.g., Bates et al.

2013; Creer et al. 2010; Lentendu et al. 2018; Mahé et al. 2017; Zinger et al. 2017). In the largest sampling of tropical sites, Mahé et al. (2017) found that hyperdiverse protist communities in lowland Neotropical rainforest soils were dominated by the parasitic Apicomplexa, and that there was low community similarity among samples collected within and between forests.

To further evaluate the biodiversity and biogeography of protist in the Neotropics, and to evaluate if the patterns found by Mahé et al. (2017) are widely applicable to non-soil environments found there, here we sampled 32 lakes associated with four different river-floodplains in Brazil. All of these lakes are often flooded by their nearby rivers, especially during the raining season. Using a metabarcoding approach with genera eukaryotic primers, we asked: 1) What are the dominate protist taxa in the Brazilian floodplain lakes?, and 2) Is there low community similarity between the Brazilian lakes?

## MATERIALS AND METHODS

### Sampling, DNA extraction, and amplification

Our study was performed in four Brazilian river-floodplain systems (**Figures S1, S2**): Upper Paraná River floodplain (Paraná, Baía, and Ivinheima rivers), Pantanal floodplain (Miranda and Paraguai rivers), Araguaia River floodplain, and Amazonian floodplain (Amazonas river). Sampling was conducted in 32 floodplain-associated lakes (**File S1**) between February and May 2012. 30 mL of water were filtered in glass fiber filters (Whatmann^®^ GF/F), and stored at a −20°C. Filters were then placed in 20 mL tubes, and DNA was extracted using Qiagen’s DNeasy Blood & Tissue kits. Extracted DNA was amplified with general eukaryotic primers that targeted the V3 hyper-variable region of the 18S rRNA locus following Nolte et al. (2010). Amplified products were then sequenced with Illumina MiSeq. The raw sequence data were deposited at ENA’s Sequence Read Archive and are publicly available under BioProject number PRJEB26716.

### Bioinformatics and statistics

All codes used here are in HTML format (**File S1**) and the OTU table is in text format (**Dataset S1**). Paired-reads were merged using VSEARCH v2.3.2 (Rognes et al. 2016) using default parameters and converted to fasta format. Assembled paired-end reads were filtered using Cutadapt v1.13 (Martin 2011) and retained if they contained both primers (minimum overlap set to 2/3 the primer length), a minimum length of 90 nucleotides, and had no ambiguous positions. Reads were dereplicated with VSEARCH and clustered using Swarm v2.1.9 (Mahé et al. 2015), with the *d* = 1 and the fastidious option on. The most abundant amplicon in each OTU was searched for chimeric sequences with VSEARCH, and their OTUs were removed. Taxonomic assignment used VSEARCH’s global pairwise alignments with the Protist Ribosomal Reference (PR^2^) database v203 (Guillou et al. 2013). Using ecoPCR v0.8.0 (Ficetola et al. 2010), the PR^2^ database was extracted for just the specific regions that were amplified and sequenced to allow for comparisons using a global pairwise alignment. Amplicons were assigned to their best hit, or co-best hits, in which case the taxonomy was resolve using the least-common ancestor with an 80% consensus threshold. The following taxonomically assigned OTUs were removed: land plants, animals, fungi, unclassified Archaeplastida, unclassified Eukaryota, and unclassified Opisthokonta. The radius of each OTU was estimated by calculating the global pairwise distance between the Swarm seed amplicon and all amplicons members of that swarm using Sumatra v.1.0.20 (Mercier et al. 2013), and by keeping the lowest similarity.

All statistical analyses were conducted in R v3.4.0 (R Core Team 2017). Dissimilarity in community compositions was analyzed based on the Hellinger transformed protist OTU matrix using the Jaccard index as implemented in the “vegdist” command in Vegan v2.4.3 (Oksanen et al. 2016). A permutational analysis of variance (PERMANOVA) was used to test for the significant effect of floodplains on this protist community dissimilarity index using the Vegan command “adonis”. A non-metric multidimensional scaling, based on the Jaccard index values, was applied to visualize the change in community composition among and between floodplains.

## RESULTS AND DISCUSSION

From a biodiversity point of view, sequencing of planktonic waters sampled from 32 lakes resulted in 1,882,036 cleaned reads and 12,081 OTUs that were taxonomically assigned to the protists (**Figure 1A**). Fewer reads and OTUs were also assigned to non-protistan taxa: 2,135,982 reads and 1,041 OTUs to the animals; 70,868 reads and 1,028 OTUs to the fungi; and 25,271 reads and 208 OTUs to the plants. About 92.3% of the protist OTUs, inferred with sequences of the V3 region, had a radius size of 98% similarity or higher (**Figures S3**). Only 0.52% of the reads and 6.17% of the OTUs assigned to the protists were <80% similar to reference sequences in the PR^2^ database (**Figure S4**); this pattern contrasts with Mahé et al. (2017), where a majority of the data from rainforest soils were <80% similar to reference sequences, but is similar to what was observed for protists from open-waters and near-shore marine environments where most of the data were similar to already-sequenced species or already-sampled OTUs (de Vargas et al. 2015; Logares et al. 2014; Mahé et al. 2017).

**Figure 1.**
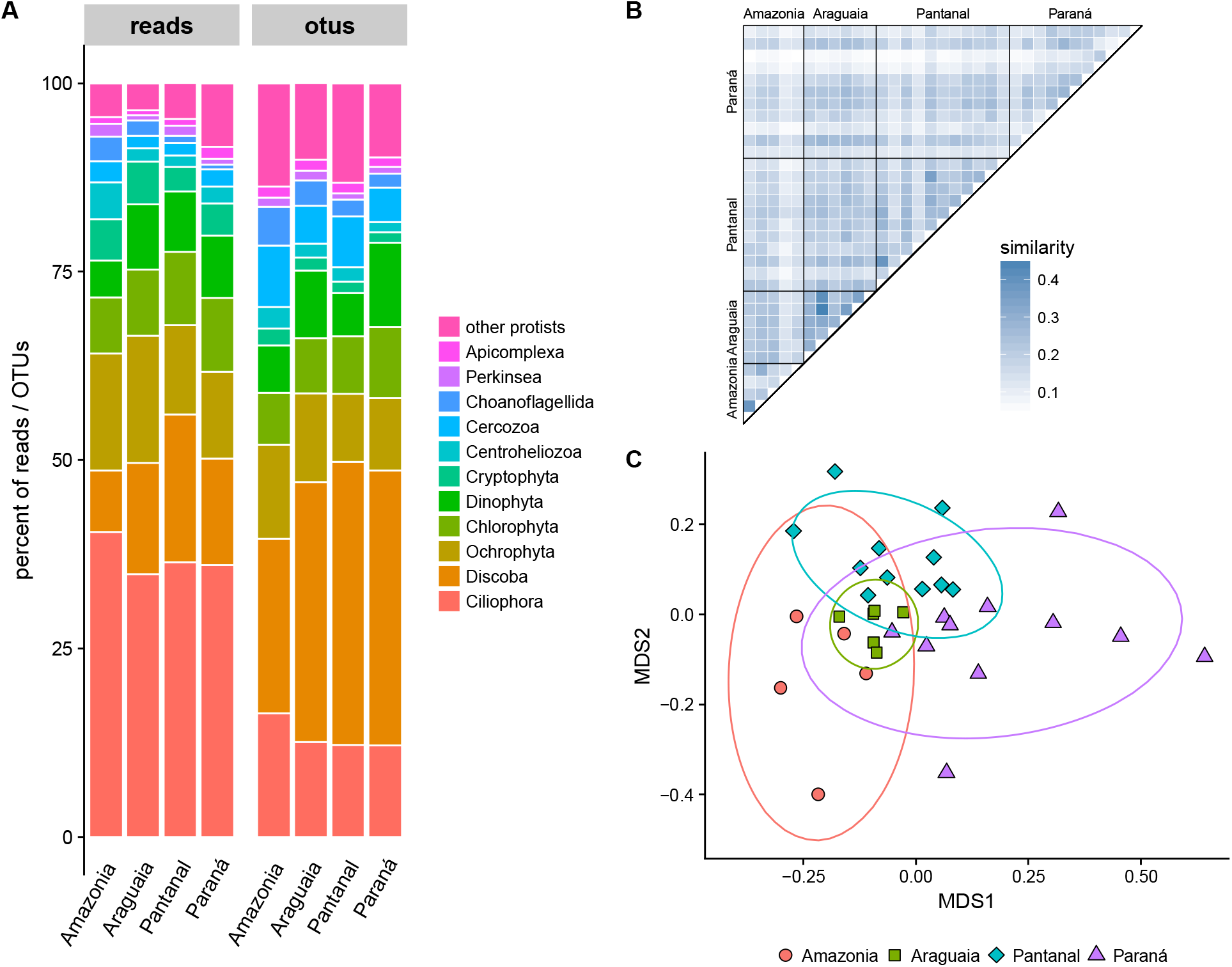
Biodiversity and biogeography of protist sampled in lakes associated with four river-floodplains in Brazil: Amazonia (Amazonas river), Araguaia River, Pantanal (Miranda and Paraguai rivers), and Upper Paraná (Paraná, Baía, and Ivinheima rivers). (**A**) Taxonomic assignment and relative abundance of reads and OTUs. (**B**) Jaccard similarity index of OTU composition differences between samples. (**C**) Non-metric multidimensional scaling of OTU composition differences between samples.

The most abundant taxon in the assigned reads was the Ciliophora, and the second was the Discoba. A slightly different pattern was found in the assigned OTUs: the Discoba was the most abundant taxon, while the second was the Ciliophora. This discrepancy between the reads and OTUs is likely explained by ciliates having highly variable differences in DNA content (Dunthorn et al. 2014; Mahé et al. 2017). This dominance of the largely free-living Discoba and Ciliophora in the protists communities from the 32 Brazilian lakes contrasts with the dominance of the parasitic Apicomplexa found in lowland rainforest soils by Mahé et al. (2017). Other groups found in high abundances in the Brazilian lakes were the largely photosynthetic Ochrophyta, Chlorophyta, Dinophyta, and Cryptophyta.

Most of the discoban OTUs were assigned to the Euglenida. This pattern of the largely photosynthetic or mixotrophic euglenids being a major component of an ecosystem has not been seen in other aquatic or terrestrial environments (e.g., Bates et al. 2013; de Vargas et al. 2015; Geisen et al. 2015; Grossmann et al. 2016; Khomich et al. 2017; Mahé et al. 2017). Why the euglenids are so abundant in these Brazilian freshwater lakes remains an ecological question; one possible explanation of this pattern is that the lakes are constantly being flooded by their nearby rivers. Euglenids can thrive in environments with high levels of humic substances and dissolved organic matter (Wetzel 2001), which is the case of these floodplain lakes where the rise in river water levels brings in decaying organic material come from macrophytes and plants into the lakes (Tockner et al. 1999). Most of these ciliate OTUs were assigned to the largely heterotrophic Oligohymenophorea and Spirotrichea. The dominance of these two ciliate groups is similar to what has been seen in aquatic ciliate communities elsewhere (e.g., Filker et al. 2016; Gimmler et al. 2016; Stoeck et al. 2014).

From a biogeographic point of view, Jaccard analyses showed low similarity in OTU compositions between all sampled lakes (mean=0.17, min=0.05, median=0.17, max=0.45; **Figure 1B**). This was the case within the floodplains (mean=0.20, min=0.05, median=0.16, max=0.28) as well as between floodplains (mean=0.16, min=0.06, median=0.20, max=0.45), although the similarity was significantly higher inside floodplains (Mann-Whitney test, p < 0.05). While low, these community similarity values where somewhat higher than what Mahé et al. (2017) found for their rainforest samples (mean=0.038, min=0, median=0.0069, max=0.96), as re-calculated by Dunthorn et al. (2017). Non-metric multidimensional scaling shows that river-floodplains cannot be clearly separated in terms of community composition (**Figure 1C**). That is, you could not predict from which river-floodplain a lake was sampled, suggesting that while community similarity was low there was also some species that were widely dispersed.

In conclusion, the patterns observed by Mahé et al. (2017) in lowland rainforest soils, in which parasites dominate communities that have low community similarity, are not necessarily applicable to other Neotropical environments. Here we found that periodically-flooded lakes associated with four river-floodplains in Brazil are dominated by the largely free-living Discoba and Ciliophora, but they community similarities between samples were likewise low. As with the diverse animal and plants inhabiting the river-floodplains, these protist communities in the lakes potentially contribute significantly to the overall diversity of protists in South American.

## ACKNOWLEDGEMENTS

Funding for this study came from the Deutsche Forschungsgemeinschaft (DFG) grant DU1319/1-1 to M.D.; Conselho Nacional de Desenvolvimento Científico e Tecnológico (CNPq) and Fundação Araucaria for all financial support of the SISBIOTA project (MCT/CNPq/MEC/CAPES/FNDCT - no 47/2010), and INPA, UNB, UFMS for infrastructure and sampling facilities to L.F.M.V.; and CNPq grant 249064/2013-8 to C.D.R.

---

**Figure S1.**
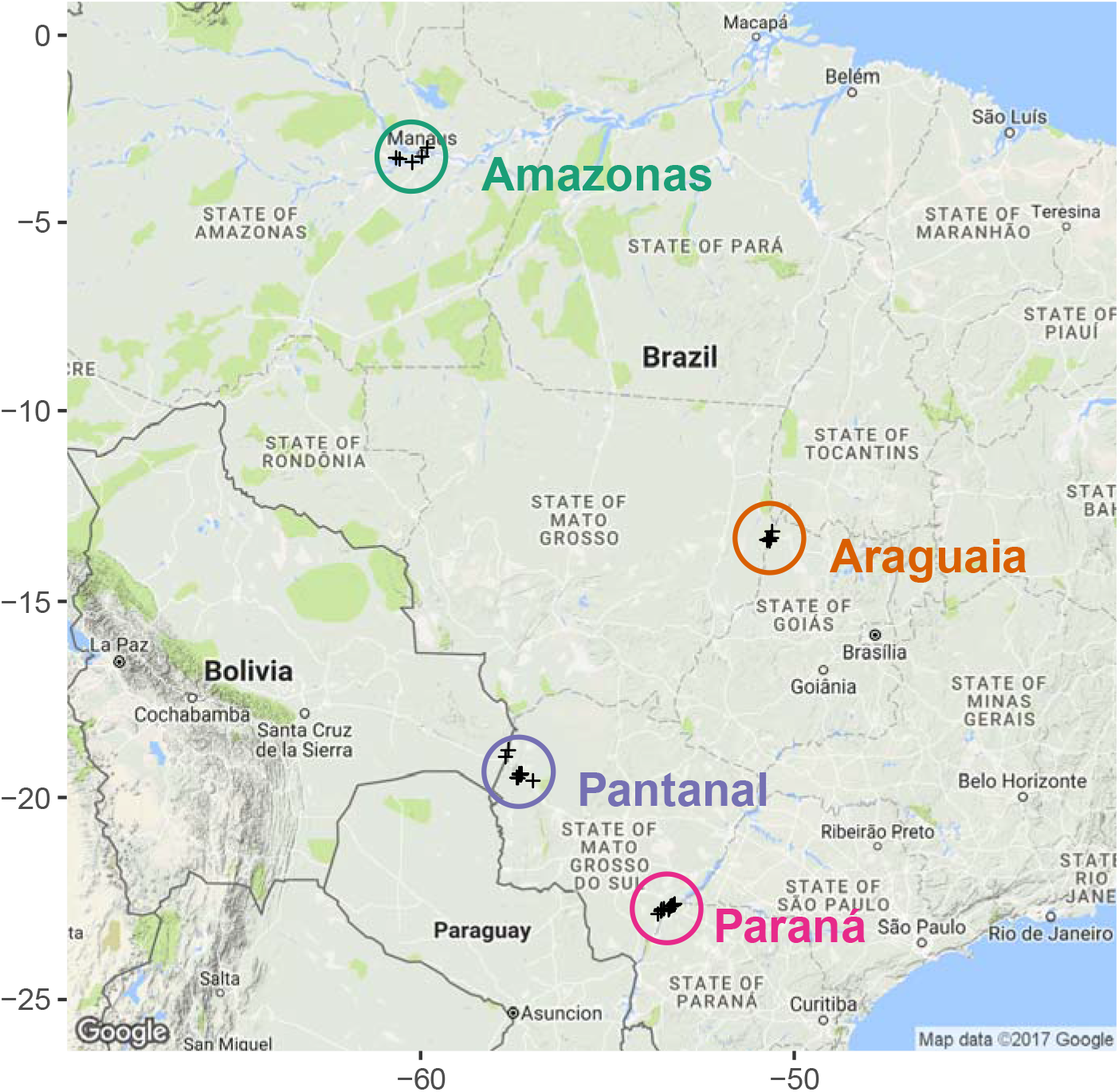
Map showing the geographic location of the four sampled four river-floodplains in Brazil.

**Figure S2.**
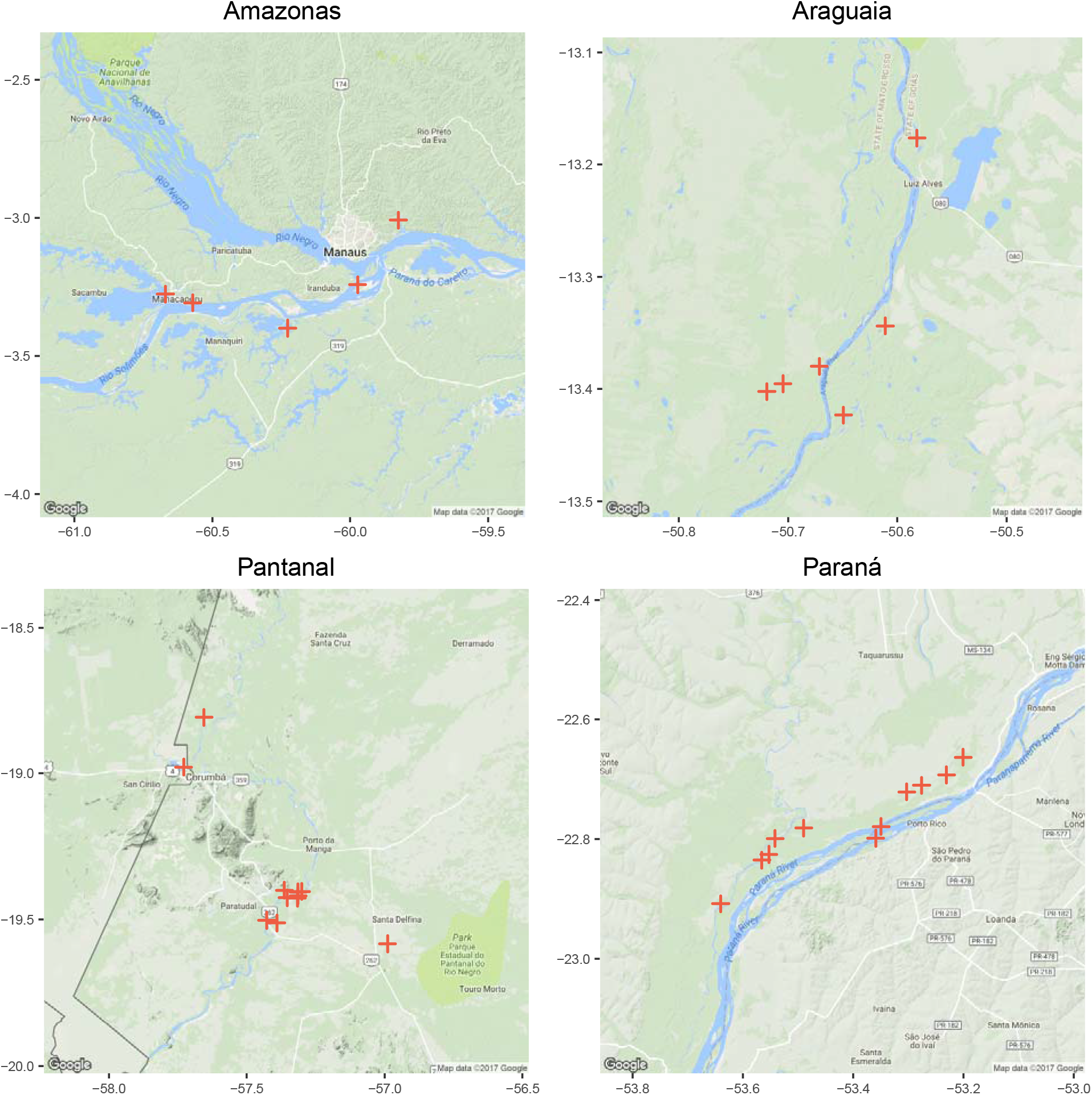
Map showing the geographic location of the sampled lakes in the different river-floodplains in Brazil.

**Figure S3.**
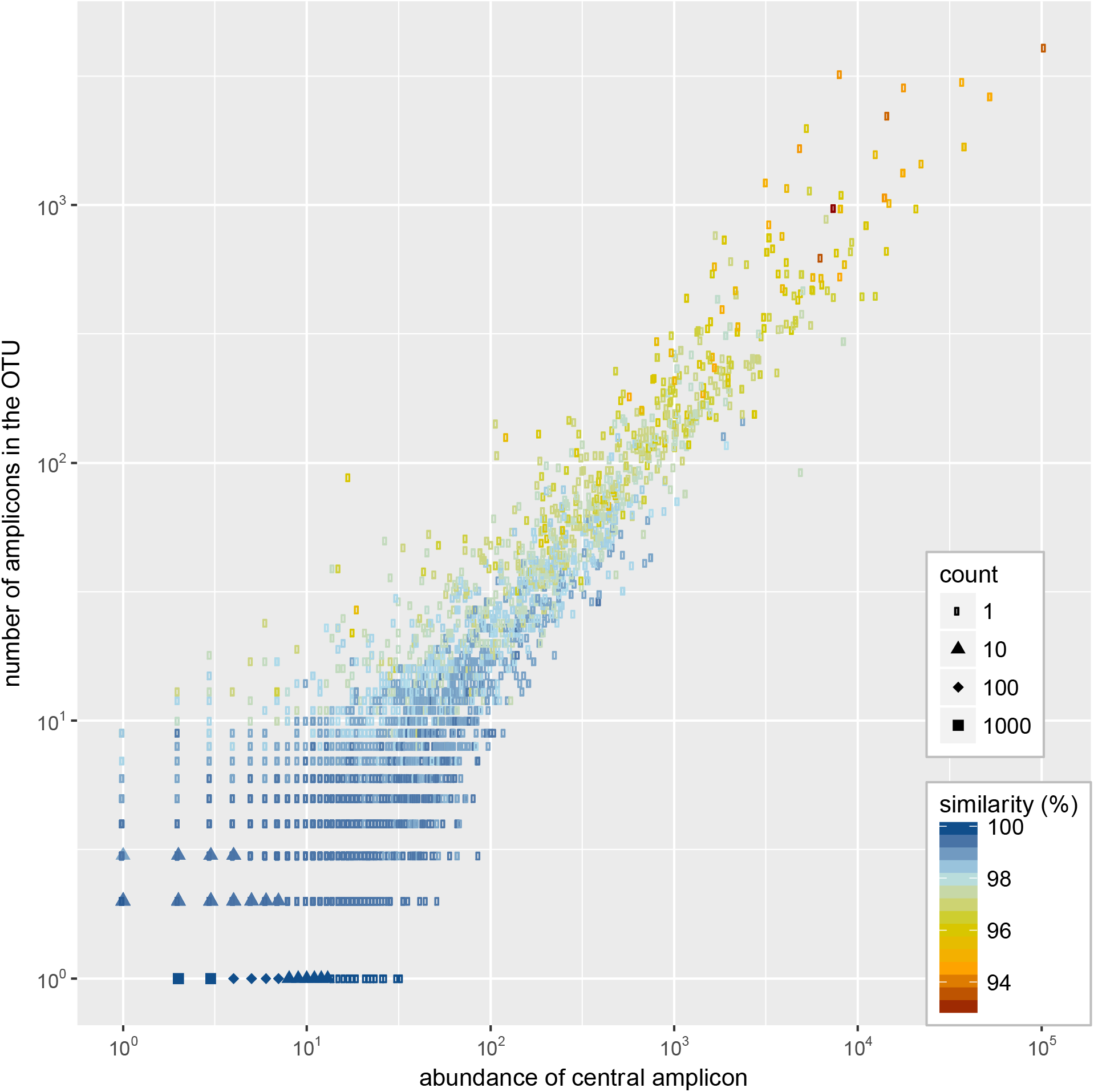
The radius (maximum % similarity between the OTU centroid and any other member of the OTU) of the protist OTUs produced by Swarm. 92.3% of the protist OTUs had a radius of 98% similarity or higher.

**Figure S4.**
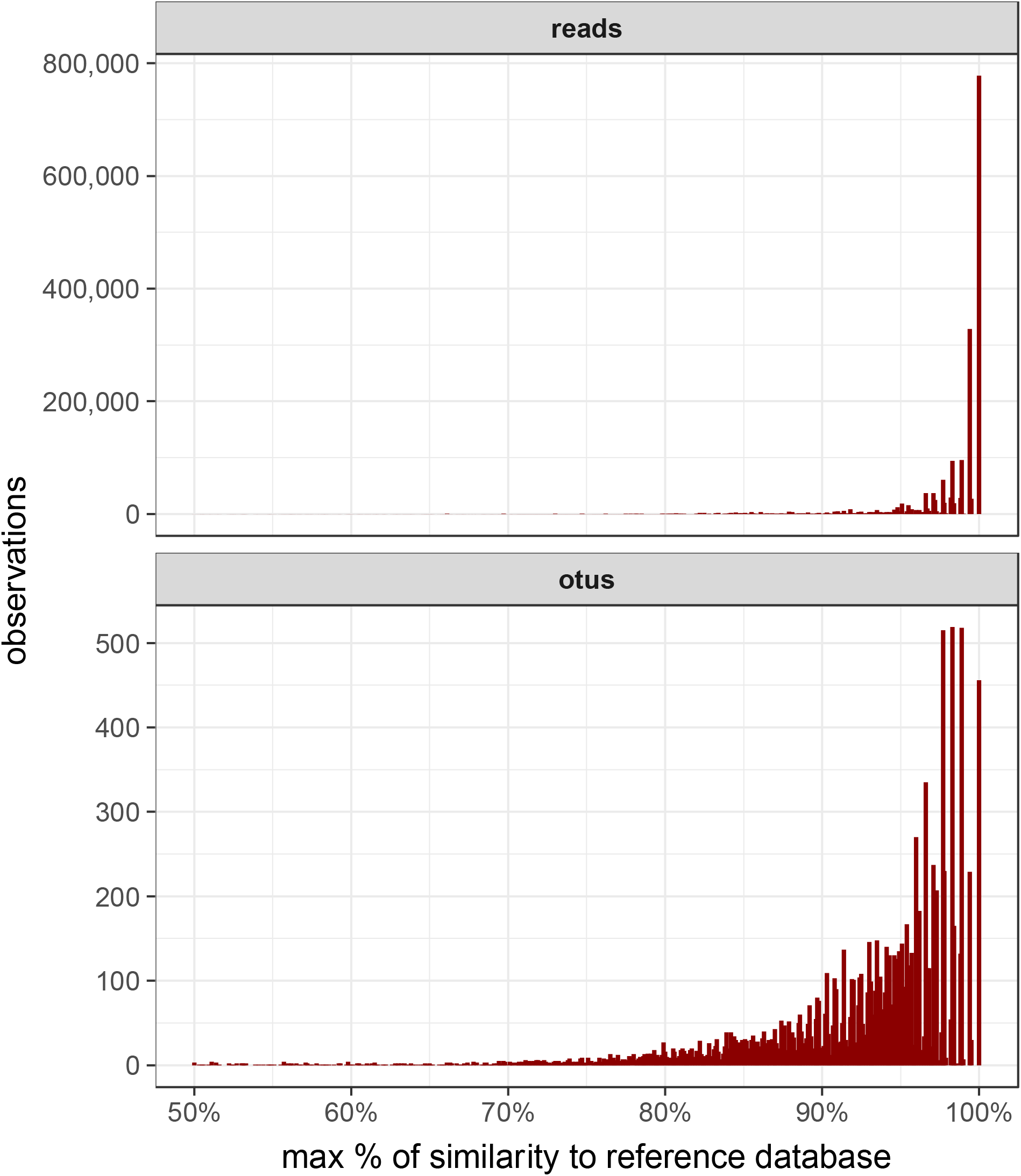
Similarity of protist reads and OTUs to the taxonomic reference database. Most of the data is >80% similar to the already sequenced species.

